# Benchmarking and integration of viral prediction tools to uncover viral dark matter across divergent low abundance metagenomes

**DOI:** 10.64898/2026.10.14.759168

**Authors:** Soufiane Alami, Grégory Destras, Maёl Rimeur, Vincent Navratil, Laurence Josset, Hussein Anani

**Author notes:** Correspondence should be addressed to Hussein Anani. Lyon 1 University, Hospices Civils de Lyon, 69004 Lyon, France., Laurence Josset. Lyon 1 University, Hospices Civils de Lyon, 69004 Lyon, France. These authors contributed equally to this work (Laurence Josset, Hussein Anani).

## Abstract

**Background:** While existing benchmark studies have focused on viral detection tools using highly abundant virus-enriched or simulated datasets, their performance in low-abundance metagenomic environments remains largely unexplored. Here, we aimed to benchmark eleven state-of-the-art viral prediction tools and assess combinations of the best-performing tools in low-abundance metagenomes.

**Methods:** We generated an artificial dataset simulating the low abundant metagenomic samples including human, bacterial and viral sequences, using complete Refseq genomes. Viral sequences were fragmented into varying lengths and subjected to increased levels of *in silico* mutations to evaluate recall, precision, and F1 score of eleven tools tested individually. We further evaluated complementary tool combinations to improve sensitivity and specificity, and assessed the best-performing multi-tools formulations on low-abundance viromes from human lower respiratory tract and air metagenomic samples.

**Results:** DeepVirfinder (DVF) and DIAMOND achieved the highest recall (median of 67% and 58%, respectively) on the RefSeq artificial contigs dataset along increased levels of induced mutations. On the artificially mutated RefSeq dataset, the highest sensitivity and F1 scores were achieved by 4 two-tools combinations : DVF+DeepMicroclass, DVF+ViraLM, DVF+geNomad and DVF+DIAMOND, combined with Kraken2X filtering to remove false positive contigs classified as Eukaryota, Bacteria, or Archaea. Using Kraken2X filtering, DVF+ViraLM and DVF+DeepMicroclass achieved the best sensitivity across real human airway and air datasets. These combinations expanded the previously identified viral repertoire using gold-standard methods, increasing the number of viral contigs with functionally annotated CDSs and revealing previously undetected viral diversity.

**Conclusion:** Benchmarking of the eleven most used tools for viral prediction showed that the complementary combination approach, particularly DVF+DMC and DVF+ViraLM coupled with Kraken2X, improves the detection of viral sequences while limiting false positives. This provides a robust strategy for uncovering divergent viral dark matter.

## Introduction

Viruses are the most abundant and diverse biological entities on Earth, with an estimated 10^7^-10^9^ distinct viral species [1]. The absence of a universal marker gene has long hindered the discovery and characterization of viral diversity compared with bacteria. However, advances in metagenomic sequencing over the past two decades have enabled the identification of vast numbers of previously unknown viruses [2]. Despite this progress, a substantial fraction of viral metagenomic sequences remains uncharacterized, constituting the viral "dark matter". Similarity-based methods such as BLAST efficiently detect viruses with limited sequence divergence from known references, as illustrated by SARS-CoV-2 discovery [3]. However, the identification of highly divergent viruses, including crAssphage, has required *de novo* assembly and Hidden Markov Models (HMM) [4]. Recent advances in viral metagenomics have driven a transition from short-read analyses to de novo assembly and viral metagenome-assembled genomes (vMAG) reconstruction [5]. *De novo* assembly followed by integrating artificial intelligence (AI)- and HMM-based tools is highly recommended nowadays for viral diversity characterization [6–8]. This enables the detection of highly divergent and emerging viral lineages that may escape gold standard methods such as Blast [9,10].

Despite these technological advancements, profiling viral communities in low-biomass settings, such as air samples, environment swabs, low respiratory tract or cerebrospinal fluids, remains largely constrained. First, current knowledge is heavily biased toward viruses from well-studied ecosystems, such as the oral and gut human virome, resulting in high levels of sequence divergence between reconstructed genomes from these samples and reference databases. Second, despite high sequencing depth, viral assemblies from low-abundance datasets are often highly fragmented, yielding viral contigs frequently shorter than 1 kilobase (Kb) [11]. This presents a real challenge for mainstream viral classification algorithms adapted to longer sequences. Most of the state-of-the-art computational predictors such as VIBRANT [11] and CheckV [12] were developed and benchmarked using high-quality and high-coverage datasets. These approaches often employ strict default filter thresholds. For example, VIBRANT requires the presence of at least 4 ORFs and >1 Kb length of viral sequences. The recent elaboration of the oral and airway virome catalog [13] used only sequences >5 Kb for viral prediction. Consequently, short fragments from low-abundant viruses are systematically discarded or misclassified, leading to a substantial underestimation of the viral diversity. Especially that ∼30% and ∼10% of reported eukaryotic and prokaryotic viruses have genome size below 5 Kb, respectively, following the latest International Committee on Taxonomy of Viruses (ICTV) report. While multiple benchmarking studies have evaluated the sensitivity and specificity of viral discovery software [14–17], these evaluations have predominantly focused on standardized, high-abundance mock communities or simulated datasets.

Despite these efforts, the performance of viral discovery tools on metagenomic data from low viral biomass samples remains insufficiently characterized. To address this gap, this study provides a comprehensive evaluation and benchmark of eleven widely used viral identification tools tested against low-abundance-like datasets.

## Methods

### Selection and configuration of viral prediction tools

Among more than 35 published tools for predicting viruses **(Table S1)** from assembled genomic and metagenomic sequences into contigs, eleven tools were selected, based on the following criteria : (i), detection capability for prokaryotic and eukaryotic viruses, and for both DNA and RNA viruses, (ii), retention of short sequences (<1 Kb), (iii), highly cited or most frequently used tools in the literature. The selected tools encompass diverse methodological frameworks, including machine-learning methods (DeepVirFinder v2020.11.21 [18] (DVF), DeepMicroClass v1.0.3 [19] (DMC), ViraLM v1.0 [20] and geNOMAD v1.11.2 [21], similarity-based methods using HMM-based (ViralVerify v1.1 [22]), k-mer-based classification (Kraken2/2X v2.1.3 [23]), or local alignment (DIAMOND v0.9.14 [24]), or hybrid approaches (Phabox2 v2.1.13 [25], CenoteTaker3 v3.4.3 [26] (CT3), CheckV v1.0.3 [12] and Virsorter2 v2.2.4 [27]). A minimum threshold of 300 bp as contig length was standardized across all benchmarking tools to maximize sensitivity for short fragments, deviating from the default settings. VirSorter2 was run with the --provirus-off option to target free viruses exclusively. For Kraken2 and Kraken2X [23], a confidence threshold of 0.1 was applied to optimize specificity, using the same custom database in both nucleotide and translated protein forms, incorporating viral taxa from RVDB (v29.0) and Inphared (v2, January 2025), as well as archaeal, bacterial, fungal, and human references from RefSeq (downloaded on 06/01/2025). DIAMOND was used in -sensitive mode to search for protein homologies using the lowest common ancestor (LCA) algorithm against the nr-NCBI database (2026). Default settings were selected to run the rest of the tools.

All tools showing a viral prediction sensitivity median above 60 % of intact sequences were selected for dual-tool combinations benchmarking, with Kraken2X or checkV used to control false positives. When Kraken2X was applied as control, all sequences classified as eukaryotic, bacterial or archaeal were filtered out. And when CheckV was selected, sequences classified as ’undetermined’ by CheckV database were excluded.

### Design of the RefSeq low abundant-like community

In order to simulate a low abundant human viral metagenome dataset, we assembled a dataset with a bacteria:human:virus ratio of 10:17:1, reflecting the metagenomic of low respiratory tract virome composition in intensive care patients as reported by Anani and colleagues [28]. This reference dataset used to construct this in silico viral metagenome comprised all complete reference genomes deposited in the RefSeq NCBI database, between January 2020 and February 2026. This includes 4,615 viral genomes, 11,260 bacterial genomes and the human genome GRCh38. Archaea and fungi were not considered because they were significantly less abundant in the respiratory tract as observed in this study [28]. To eliminate viruses integrated into bacterial genomes (prophages), a rigorous two-step cleaning process was applied: (i) identification and extraction of integrated prophages using Phigaro (v2.4.0) [29]. (ii) exclusion of any bacterial sequence in which more than 30 % of the ORFs showed homology with viruses from the pVOG database [30], following the strategy described in the study of Ho and colleagues [31]. Finally, all viral, bacterial and human genomes were dereplicated at ≥95 % nucleotide identity and ≥85 % coverage using CD-HIT v4.8.1 [32] and fragmented using custom Python script into contigs/fragments with lengths ranging from 300 to 2,000 bp to mimic the contig sizes typically obtained in respiratory origin contigs reconstructed from Illumina sequencing data. Random subsampling using seqkit generated ten distinct (non-overlapping) *in silico* replicates adhering to the 10:17:1 ratio (Bacteria: Human: Virus) and yielding 100,000 bacterial fragments, 170,000 human fragments, and 10,000 viral fragments per replicate (corresponding to 4,615 viral genomes).

### Generation of divergent viral fragments by *in silico* mutagenesis

To assess the impact of genetic divergence on the performance of prediction tools, the viral fragments from the ten replicates underwent an *in silico* mutagenesis using the mutate.sh function of the BBTools suite v39.26 [33]. Ten levels of divergence were generated, ranging from 95% to 50% nucleotide identity compared with the original intact sequences serving as the 100% reference. The induced mutations include insertions (0.03%), deletions (0.03%), and, predominantly, substitutions (4.94%) at each 5% divergence step. Sequences with up to 50% divergence were generated by progressively introducing mutations.

### Performance evaluation metrics

We evaluated the prediction performance of the used tools to distinguish viral sequences from non-viral sequences. The predictions were compared with the original Refseq viral genomes to classify intact or mutated viral fragments into the following categories : True Positive (TP) or False Negative (FN) for viral sequences, and True Negative (TN) orFalse Positive (FP) for non-virals sequences. These categories were defined as the following : True Positive (TP) : refers to viral contigs that are identified as viruses, False Negative (FN) : refers to viral contigs identified as non-viral, False Positive (FP) : these are non-viral contigs classified as viruses, True Negative (TN) : these are non-viral contigs identified as non-viral. Then, we calculated performance metrics of each tool as previously described [15,34,35]. Sensitivity measures the ability to correctly identify viral contigs, while precision evaluates the proportion of truly viral contigs among the predicted viral contigs. Specificity assesses the correct exclusion of non-viral contigs. Finally, the F1 score, the harmonic mean of the sensitivity and precision, provides an overall measure of the prediction performance. These metrics are calculated using the following equations :

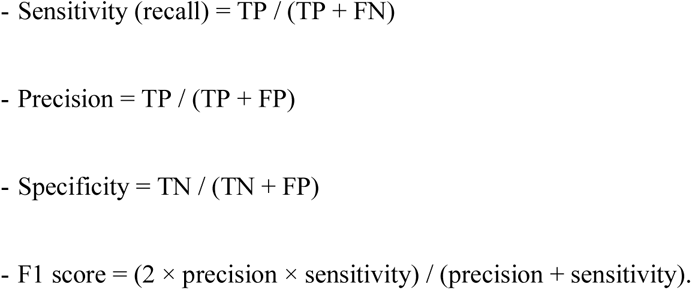

### Reanalysis of human low respiratory tract viral metagenomes

A total of 121 endotracheal samples collected from 87 critically ill patients admitted to the intensive care unit were retrieved from a clinical cohort (IBIS) and sequenced using viral metagenomics approach. DNA and RNA extracted from the endotracheal aspirates were sequenced using a viral metagenomic approach. Double and single stranded DNA and RNA viruses can thus be retrieved. The dataset was obtained from the European Nucleotide Archive (ENA) under BioProject accession PRJNA1110359. These samples were previously shown to contain a diverse range of prokaryotic and eukaryotic viruses, as reported by Anani and colleagues [28]. The sequencing reads were processed using the same bioinformatic pipeline described in the original study, resulting in the generation of 256,304 dereplicated contigs. All evaluated viral identification tools were subsequently applied directly to these contigs using the same parameters as those used for the in silico datasets. A minimum contig length threshold of 450 bp was applied, consistent with the criteria defined in the original study. The reference viral set was identified using Diamond similarity searches, following the methodology and classification criteria established by Anani et al. These viral contigs (26,822) were considered as the reference dataset (ground truth) for evaluating the performance of the tested viral detection approaches.

### Reanalysis of environmental (air) viral metagenomes

To evaluate the performance of viral identification tools on environmental samples containing low-abundance viruses, we downloaded the raw sequencing data corresponding to 13 air metagenomic samples from the NCBI Bioproject PRJNA854686 and generated by Deng and colleagues [36]. Air samples were collected in Beijing, China, using the Bioaerosols and Dust Sampling Payload, under five different atmospheric conditions : sunny, cloudy, rainy, lightly and moderately polluted, with three biological replicates obtained for each condition. DNA extracted from the collected bioaerosols was sequenced using a shotgun metagenomic approach. In this dataset, only double and single stranded DNA viruses can thus be retrieved. To ensure consistency with the original study, the raw sequencing reads were processed using the same bioinformatic workflow described by the authors to generate *de novo* assembled, dereplicated contigs. First, this workflow initially identified 13,147,416 instead of 16,021,599 contigs as reported in the original study, with a minimum contig length of 300 bp. Second, subsequent size filtering (not shorter than 300 bp) and quality assessment and filtering using a combination of checkV and VirSorter2 resulted in the identification of 141,506 of the reported 141,576 high-confidence viral contigs. These viral contigs were used as the reference set (ground truth) for evaluating the performance of the tested viral detection approaches.

### Functional annotation and sequence compositional analysis

Functional annotation was performed for all viral contigs corresponding to the reference of the respiratory and air datasets, including those predicted as viral from all tool combinations. Viral contig encoded CDSs were predicted using Prodigal v2.6.3 [37]. Predicted viral contigs were subsequently annotated using the Pharokka v1.10, Phold v1.2.6, Phynteny transformer v0.1.3 workflow [38], which integrates protein sequence homology, structural similarity, and gene synteny to maximize viral protein annotation and functional assignment.

Remaining unannotated contigs were compared against the GRCh38 human reference genome using BLASTn (using the threshold identity of 85% and alignment length of 50 bp) and representative bacterial genomes using GTDB-Tk v2.7.2 [39] (with default parameters). To evaluate the quality of unannotated viral contigs and identify potential chimeric sequences, we performed a compositional analysis based on tetranucleotide frequency (TNF) and G+C content. For each viral contig, we calculated overall sequence size and G+C content using the R library Biostrings v.2.76.0. Tetranucleotide frequencies [40] were computed as the relative frequencies of all 256 possible four nucleotide oligomers (or 4-mers) along each contig. To assess the intra-contig compositional homogeneity, each contig was divided into sliding windows of 300 bp with a 100 bp step size, and G+C content and TNF were recalculated at each window. For each contig, we then quantified compositional variability across windows using two metrics: (i) the G+C range, defined as the difference between the maximum and minimum window-level G+C content, and (ii) the mean tetranucleotide standard deviation (“TNF_mean_sd”), calculated as the average, across all 256 tetranucleotide frequencies, of the standard deviation observed across sliding windows. Contigs exhibiting a G+C range greater than 15 % and/or a mean TNF standard deviation greater than 0.02 were flagged as "potential chimeras," reflecting a shift in nucleotide composition indicative of a putative assembly artifact joining sequence fragments from distinct source organisms. Contigs not meeting either threshold were classified as "compositionally consistent".

## Results

### Viral prediction tools show variable performance on simulated respiratory-like divergent viral metagenomes

In order to evaluate the robustness of viral identification tools applied to low viral abundance and increased sequence divergence, we generated an *in-silico* dataset comprising viral, bacterial and human contigs **(Figure 1a)**. The proportions of each sequence type were designed to mimic the microbial composition of the low abundance environment such as low respiratory tract metagenomes, in line with the bacteria:human:virus ratio of 10:17:1 as reported by Anani and colleagues [28]. After dereplication, the viral RefSeq dataset comprised 4,615 viral genomes. Of these, 68% were prokaryotic viruses, including 94% of DNA and 6% of RNA phages. The remaining 32% were eukaryotic viruses, comprising 73% of RNA and 27% DNA viruses **(Figure 1b)**. Among DNA viruses, *Caudoviricetes* phages were the most represented group (n=2501/3132). Among RNA viruses, *Bunyaviricetes* (397/1483), *Monjiviricetes* (246/1483), and *Pisoniviricetes* (148/1483) were among the most represented classes **(Figure 1c)**.

**Figure 1:**
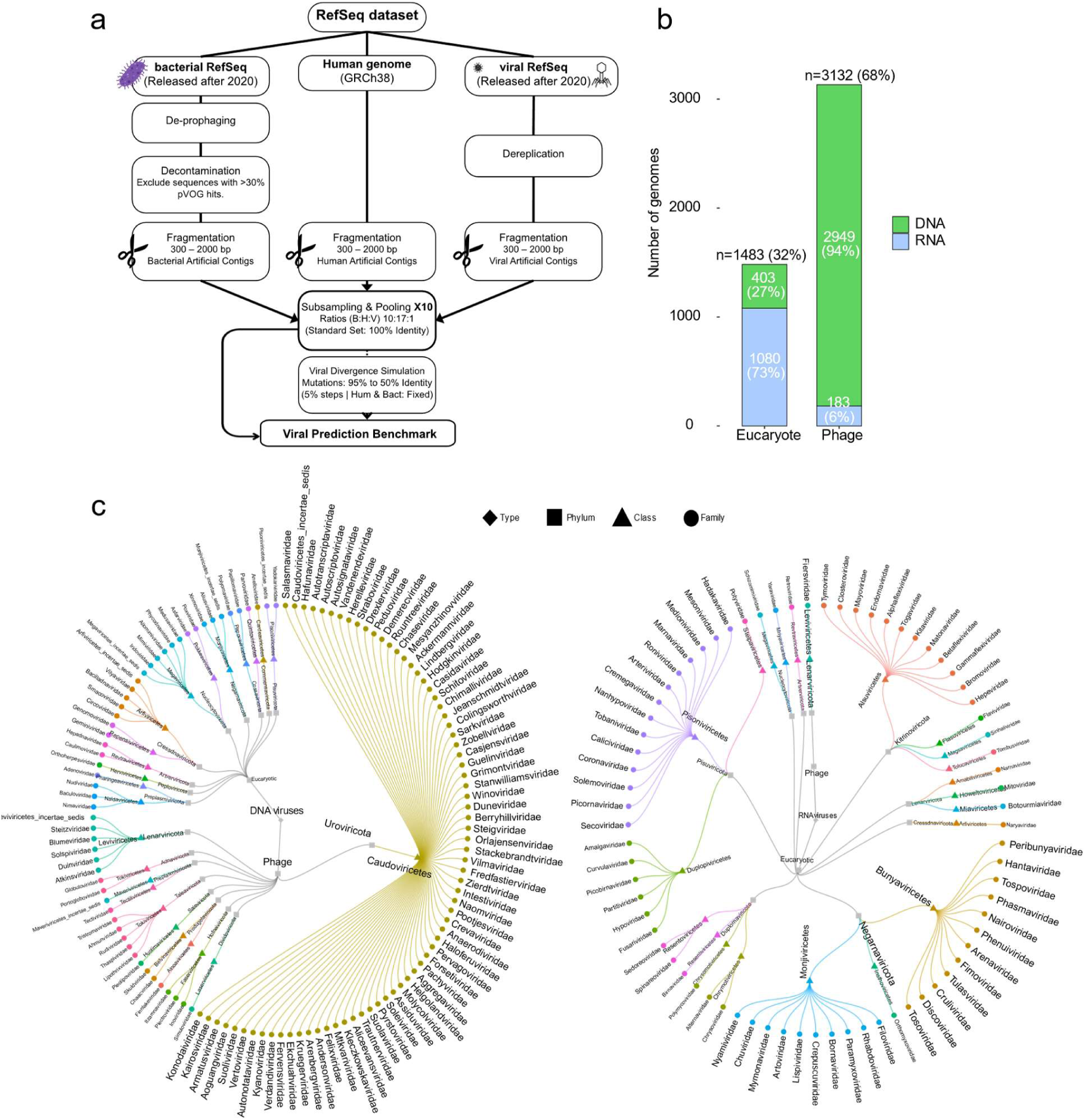
Overview of the benchmarking workflow. a, Schematic representation of the workflow used to construct the simulated respiratory-like viral RefSeq dataset for benchmarking viral prediction tools. b, Distribution of host origins and virus types represented in the RefSeq dataset. c, Dendrogram depicting the viral diversity represented in the dataset at the family, class, and phylum levels.

After fragmenting RefSeq sequences, the dataset was partitioned into ten independent, non-overlapping replicates, each containing ∼10,000 viral contigs. Viral sequences were then computationally mutated to generate divergence levels ranging from 0% to 50% relative to the original intact sequences. We evaluated eleven state-of-the-art viral identification tools (DVF, DMC, ViraLM, geNomad, ViralVerify, Kraken2/2X, DIAMOND, Phabox2, CT3, CheckV and Virsorter2) across the resulting benchmark datasets **(Table S1)**.

On the whole unmodified datasets (0% divergence), the tools achieving the highest recall were Kraken2X (99.8% recall), ViraLM (93%), DVF (85.3%), geNomad (86%), Kraken2 (83%), DMC (78.8%) and DIAMOND (66.2%), demonstrating excellent sensitivity in the detection of known viral sequences **(Figure 2a, Table S2** and **Supplementary Data 1**). However, recall was below 60% for Phabox2, VirSorter2, CT3 and ViralVerify. When predictions were stratified by sequence size, Kraken2X achieved the highest recall (∼100%) consistently across all contig size categories (**Supplementary Figure 1a, Table S3**). Phabox2 and VirSorter2 recall increased with sequence length from 1,000 bp and 1,500 bp, respectively, whereas DMC demonstrated consistent recall efficiency (>80%) for sequences of at least 500 bp as it does not predict sequences shorter **(Supplementary Figure 1a).**

**Figure 2:**
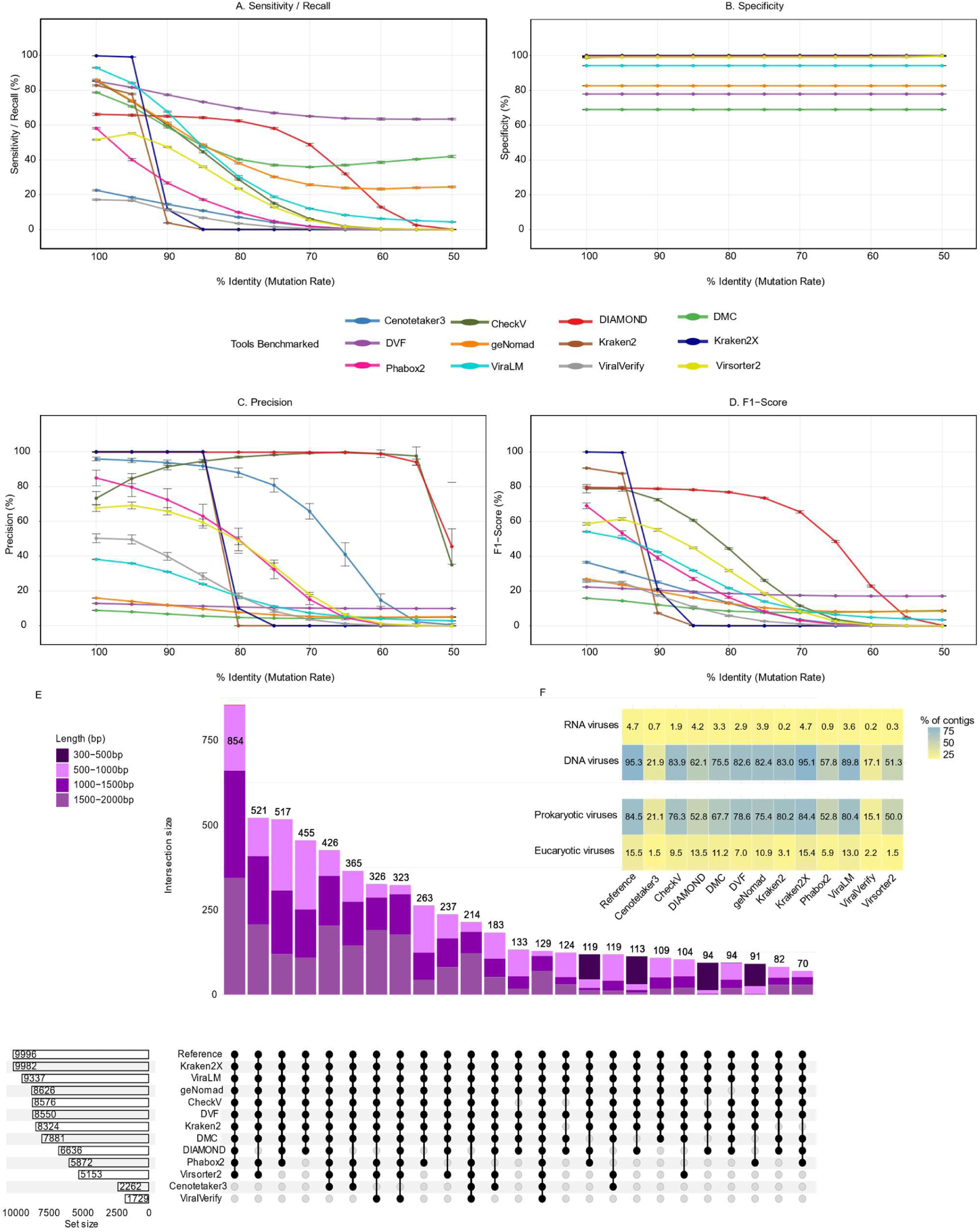
Viral prediction performance of the eleven evaluated tools on simulated respiratory-like divergent viral metagenomes. a-d, Sensitivity (recall) (a), specificity (b), precision (c), and F1 score (d) of the evaluated tools on intact and mutated RefSeq viral sequences. (e) UpSet plot showing the intersections of predicted viral contigs across all tools using a RefSeq simulation dataset comprising 9,996 viral contigs. (f) Distribution of viral predictions across virus host origins and virus types.

DMC, DVF, geNomad, and ViraLM showed reduced specificity and higher false-positive rates compared to other predictors **(Figure 2b).** Precision also varied considerably among the tested tools, ranging from 10% with DMC to 100% with Kraken2, Kraken2X and DIAMOND **(Figure 2c)**. The precision of Kraken2X, Kraken2, DIAMOND and CT3 was consistently high across all contig length categories, exceeding 90% in each case **(Supplementary Figure 1b)**. Finally, Kraken2X, Kraken2, DIAMOND and Phabox2 showed the highest F1 score with 100%, 90.6%, 79.5% and 69%, respectively **(Figure 2d).**

When benchmarking tools with *in silico* mutated viral sequences, Kraken2 and Kraken2X performed poorly in predicting viruses in sequences with 10% divergence or more, showing a steep decline in recall, precision and F-score **(Figure 2a-d, Supplementary Data 1)**. DVF maintained a stable recall until 50% sequence divergence and outperformed the other tools with a sensitivity rate ranging from 85.3% to 63.5%. Among the remaining tools, performance varied substantially. Diamond maintained an acceptable level of recall up to 30% sequence divergence, whereas ViralVerify and CT3 performed poorly across all divergence levels **(Figure 2a)**. At the precision level, DIAMOND performed well with a stable profile over the divergence rate. The other tools performed differently : CT3 showed good performance up to 20% divergence, then started to decline ; DVF, geNomad and DMC were inaccurate in this context **(Figure 2c)**. Finally, looking at the F1 score, we observed a stable performance of DIAMOND despite increased divergence in viral sequences (Mean F1 score ∼ 78%, **Figure 2d**). Across all contig size classes, DVF achieved the highest mean recall, with values of 79.59%, 87.54%, 84.08% and 86.58% for the 300-500 bp, 500-1000 bp, 1000-1500 bp and 1500-2000 bp intervals, respectively **(Supplementary Figure 1a)**. This was consistent for F1 score in the 500-1000 window, however DIAMOND showed the best performance at all and especially with sequence divergence lower than 30% **(Supplementary Figure 1a-c)**.

Among all evaluated tools, DIAMOND was associated with the most important computational time constraint (∼47 hours per CPU to predict 1,000 contigs with size ranging between 300 bp and 2,000 bp) compared with the other tools (in median 0.2 hours per CPU to predict 1,000 contigs) **(Supplementary Figure 2a** and **Table S4).**

Finally, when we compared the predicted viral contigs of all the tools studied on a simulation of intact sequences (9,996 RefSeq viral contigs), strikingly, we found that only ∼8% of the contigs overlapped between all the tools **(Figure 2e)**. This was consistent with the ability of each tool to predict viral types and host origin, with Kraken2X showing the highest prediction performance, followed by ViraLM and Kraken2, across DNA, RNA, prokaryotic, and eukaryotic viruses **(Figure 2f)**. These findings highlight differences in the performance of the underlying algorithms for viral prediction.

### Combining viral prediction tools improve virus detection on simulated respiratory-like divergent viral metagenomes

We then investigated whether combining tools could improve recall, precision and the F1 score. We evaluated all possible combinations of the five tools achieving >60% recall on intact sequences (0% induced mutations), resulting in ten dual-tool combinations. To control false-positive rates, each tool combination was subjected to an additional filtering step using either Kraken2X or CheckV. For Kraken2X, sequences classified as eukaryotic, bacterial, or archaeal were removed, whereas sequences classified as “undetermined” by CheckV were excluded.

Combined with Kraken2X filtering, four dual combinations (DVF+DIAMOND, DVF+DMC, DVF+geNomad and DVF+ViraLM, **Supplementary Figure 2** and **Table S4**) showed the best stable recall profile (mean recall of 82%, 80%, 78% and 76%, respectively) up to 50% divergence rate (**Figure 3a** and **Table S5**). This profile was conserved across all contig size windows, where DVF+DIAMOND showed the best performance across all size fragments until 30% of divergence rate, with a mean of 72.2%, 86.9%, 80.7% and 82.9% at 300-500, 500-1000, 1000-1500 and 1500-2000 bp **(Supplementary Figure 3a, Table S5)**. When taking into account the whole divergence scale, DVF+DMC showed the best recall with a mean score of 86.3%, 79.6% and 80.6% at 500-1000 ; 1000-1500 and 1500-2000 bp length ranges **(Supplementary Figure 3a)**. Importantly, these scores were systematically above the recall scores of DVF alone, the tool with the highest performances for highly divergent viral genomes. Finally, DVF+geNOMAD and DVF+ViraLM showed the best recall for very short contigs (300-500bp) with a mean of 70.1% and 66.5%, respectively **(Supplementary Figure 3a)**. The performance of these four combinations (DVF+DIAMOND, DVF+DMC, DVF+ViraLM and DVF+geNomad) was conserved at the F1 score level with a mean score of 82.1%, 79.5%, 75.9% and 78.2%, respectively over all divergence levels **(Figure 3a)**. Again, these scores were higher than those observed with DIAMOND or DVF alone, the tools with the highest F1 score when tested alone, with 30% and 50% of divergence rate respectively. After Checkv filtering, all combinations exhibited a significant drop in performance in terms of either recall or F1 score at a divergence rate of 0 to 50% **(Figure 3b** and **Supplementary Figure 3b**).

**Figure 3:**
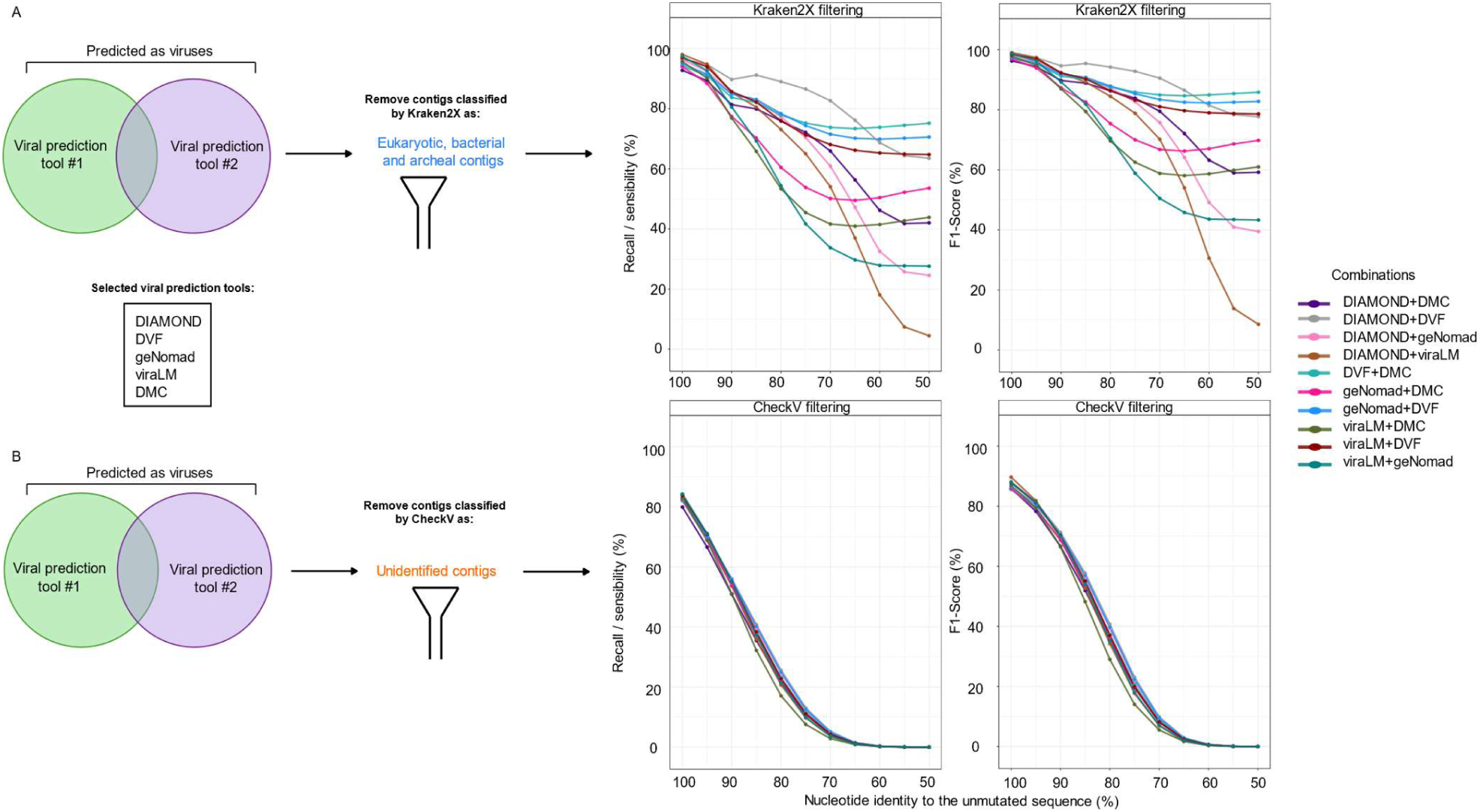
Viral prediction performance of the tool combinations on simulated respiratory-like divergent viral metagenomes. a-b, Sensitivity (recall) and F1 score metrics of the evaluated tool combinations using Kraken2X (a) and CheckV (b) for false positives filtering.

Finally, with 50 and 48 h/CPU respectively, DIAMOND+DVF+CheckV and DIAMOND+DVF+Kraken2X required the highest computation time, compared with other combinations that had a median of 2.6 h/CPU (**Supplementary Figure 2b** and **Table S4**). These results suggest that combining performance of different tools can enhance overall viral prediction on divergent viral genomes.

### Combinations of viral prediction tools reduce the viral dark matter of the lung virome

We next evaluated the performance of the best-performing tool combinations on real viral metagenomic data generated from the lower airway fluids of critically ill patients [28]. Raw sequencing data were obtained from the original study and processed using the proposed pipeline to reproduce the assembly of viral contigs. As described previously, all putative false-positive contigs were discarded using Kraken2X or CheckV, therefore, only the sensitivity of each combination was evaluated. Among the 256,304 assembled contigs, 26,822 viral contigs were identified in the original study and considered the reference viral set. Applying the four selected combinations (DVF+DMC, DVF+ViraLM, DVF+geNOMAD, and DVF+DIAMOND) coupled with Kraken2X filtering yielded 60,217, 59,614, 25,917, and 23,952 predicted viral contigs, respectively, compared to 26,863, 33,682, 21,365, and 16,867 after CheckV filtering **(Figure 4a-b**, **Table S6** and **Supplementary Data 2)**. The intersection of the four combinations predictions with the reference, showed a recovery of ∼72% (n=19233/26822) and ∼53% (n=14179/26822) of the reference dataset when filtering false positives with Kraken2X and checkV, respectively **(Figure 4b,c)**. 24,054 (DVF+DMC), 29,792 (DVF+ViraLM), 19,115 (DVF+geNomad) and 15,921 (DVF+DIAMOND) of contigs identified by Kraken2X combination were shared with those filtered by CheckV. To investigate whether additional viral sequences identified by the combinations carried different viral functionals, we screened for putative viral coding genes. Annotation of the viral contigs showed that nearly 100% of the identified sequences contained viral coding genes, with a profile comparable to that observed in the reference **(Figure 4d, Supplementary Data 3)**. The functional composition of the combinations viral contigs was similar with that of the reference across all tool combinations **(Figure 4e)**. However, the DNA, RNA and nucleotide metabolism functions were more abundant (Fisher test p < 0.00001) in the combination DVF+DMC+Kraken2X compared to the reference.

**Figure 4:**
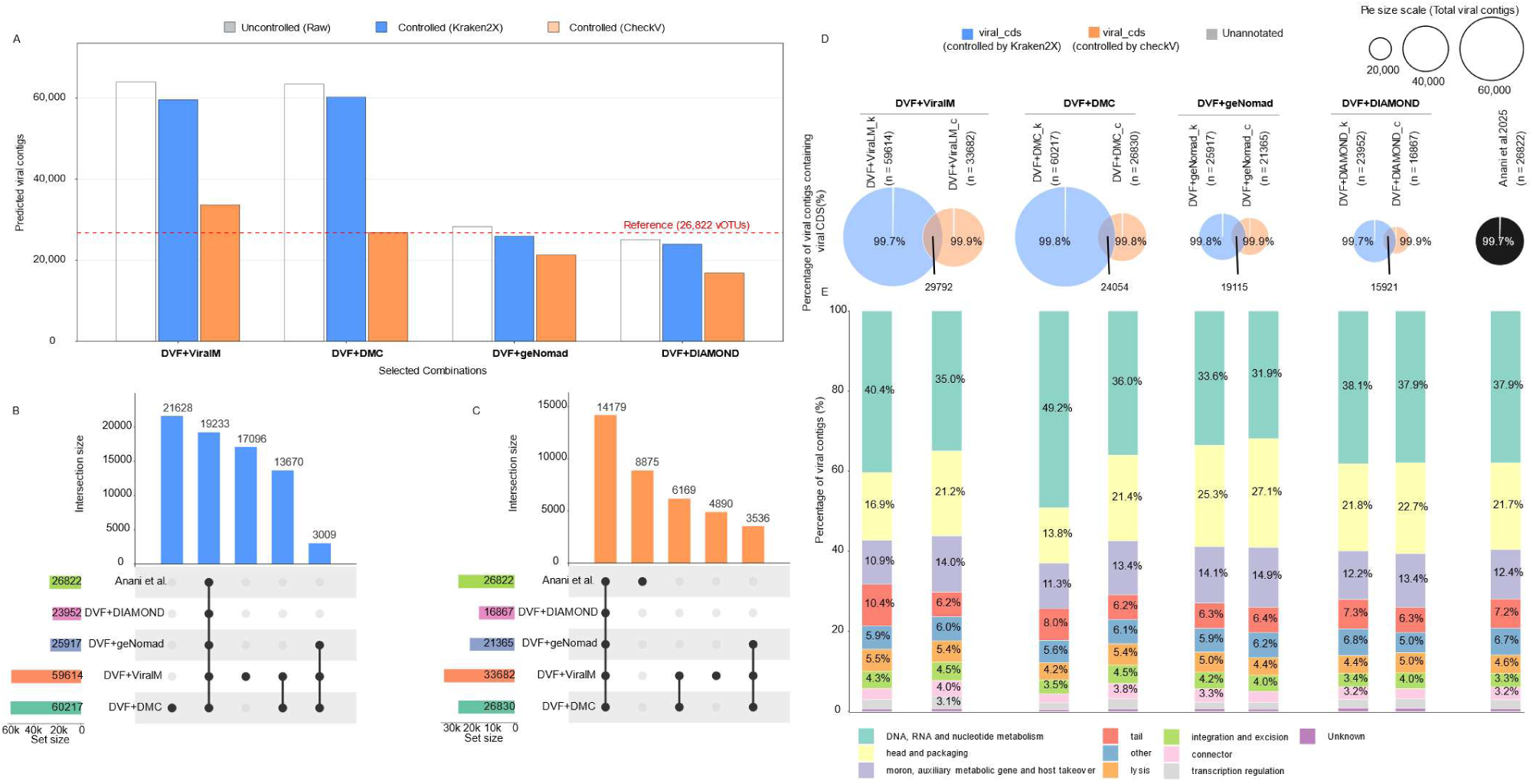
Application of the four best-performing tool combinations reduces viral dark matter in the lung virome. a, Bar plot showing the number of predicted viral contigs before and after Kraken2X and CheckV filtering for the four best-performing tool combinations. b-c, UpSet plots showing the intersections of predicted viral contigs across all tool combinations after Kraken2X (b) and CheckV (c) filtering. d, Bubble plot showing the percentage of viral contigs containing viral CDSs across all evaluated combinations compared with the reference dataset. Combinations with Kraken2X (_k) filtering are colored in blue and those with CheckV (_c) in orange. e, Functional profiles of predicted viral contigs recovered by each tool combination.

These results emphasize the potential of combining viral prediction tools to increase the capacity for discovering viral sequences.

### Combinations of viral prediction tools reduce the viral dark matter of airborne virome

We then assessed the performance of the four combinations on the viral airborne metagenomic data [36]. Raw data from 13 air samples collected at an altitude of 55 m in China, was downloaded and manipulated using the methodology steps provided in the study in order to obtain the viral contigs. Of the 141,567 high-confidence viral contigs reported in the original study, we successfully reproduced the proposed pipeline and recovered 141,506 high-confidence viral contigs out of 13,147,416 assembled contigs, that will be used as the airborne reference for benchmarking. Applying the four selected combinations (DVF+DMC, DVF+ViraLM, DVF+geNOMAD, and DVF+DIAMOND) coupled with Kraken2X filtering, resulted in 217,045, 162,480, 143,434, and 142,930 viral contigs, respectively, compared to 66,874, 55,824, 45,699 and 44,160 after CheckV filtering **(Figure 5a-b, Table S7** and **Supplementary Data 4**). Importantly, the intersections of the combinations with Kraken2X or checkV filtering recovered only ∼2% and ∼28% of the reference dataset, respectively **(Figure 5b,c**). Only 4,046 (DVF+ViraLM), 4,015 (DVF+DMC), 3,402 (DVF+geNomad) and 3,021 (DVF+DIAMOND) of contigs identified by Kraken2X combination were shared with those filtered by CheckV **(Figure 5d).** The annotation of the viral contigs showed that after Kraken2X filtering, 89.1%, 88%, 83.4% and 73.4% in DVF+geNomad, DVF+ViraLM, DVF+DMC and DVF+DIAMOND, respectively, harbored viral coding genes. After CheckV filtering, nearly 100% of the newly identified contigs harbored viral coding genes **(Figure 5d** and **Supplementary Data 5)**. Finally, the functional profiles of viral contigs in each combination after CheckV filtering were similar to the reference with the DNA, RNA and nucleotide metabolism as the most abundant function (p<0.0001). In contrast, Kraken2X filtered contigs showed distinct functional schemes with an enriched presence of the moron, auxiliary metabolic gene and host takeover (p<0.0001; **Figure 5e** and **Supplementary Data 5).** Most of the contigs with no known viral coding genes (>99.9%) contained neither human nor bacterial coding genes (**Supplementary Figure 4a-b** and **Supplementary Data 6**). When we analysed their compositional structure, we found that 13,976 unannotated viral contigs were shared across all combinations (**Supplementary Figure 4a**). Of these contigs, only ∼0.05% aligned to the human genome (**Supplementary Figure 4b**). Their lengths ranged from 0.3 to 30 Kb, with a G+C content ranging from near 20% to 60% (**Supplementary Figure 4c-d)**. Tetranucleotide frequency analysis (TNF) revealed that a small proportion of contigs exhibited heterogeneous nucleotide composition and G+C content, indicating the presence of <10% of potential chimeric sequences. Overall, >90% of the newly identified contigs exhibited consistent compositional profiles (**Supplementary Figure 4f-h** and **Supplementary Data 7)**.

**Figure 5:**
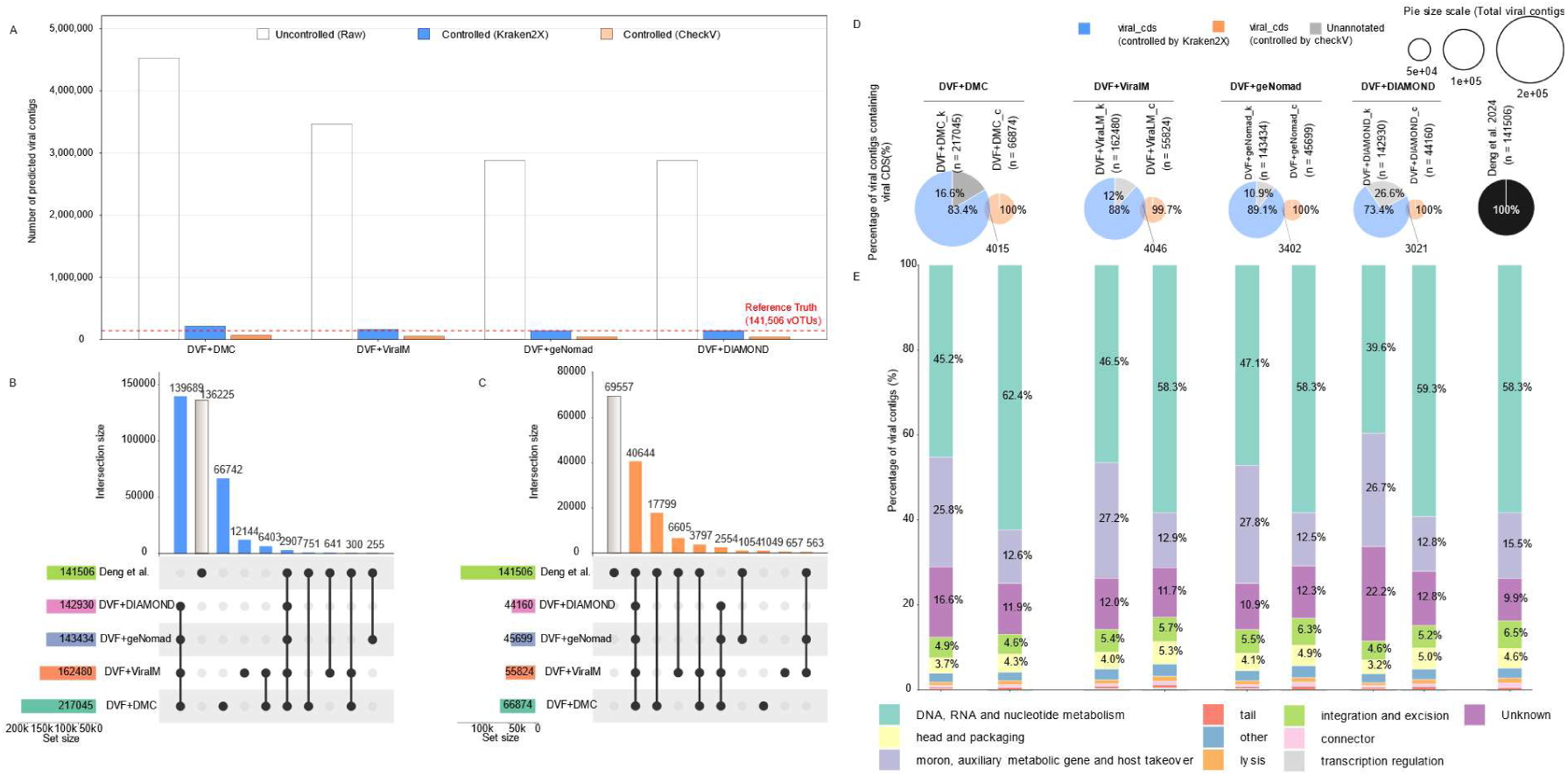
Application of the four best-performing tool combinations reduces viral dark matter in the airborne virome. a, Bar plot showing the number of predicted viral contigs before and after Kraken2X and CheckV filtering for the four best-performing tool combinations. b-c, UpSet plots showing the intersections of predicted viral contigs across all tool combinations after Kraken2X (b) and CheckV (c) filtering. d, Bubble plot showing the percentage of viral contigs containing viral CDSs across all evaluated combinations compared with the reference dataset. Combinations with Kraken2X (_k) filtering are colored in blue and those with CheckV (_c) in orange. e, Functional profiles of predicted viral contigs recovered by each tool combination.

These results confirm the high capacity of the proposed combinations to annotate previously undescribed viral sequences.

## Discussion

In this study, we have pinpointed the performance of viral detection tools in environments with low virome abundance, where viral metagenomic sequences are often obscured by abundant microbial and host-derived nucleic acids. Our findings revealed significant variations in tool performance across datasets, with no method achieving consistent results on all metrics levels. Therefore, we investigated the combinations of viral prediction tools and demonstrated that integrating complementary approaches can improve predictive performance. We have shown that four combinations, each including DVF and either another deep-learning tool (DMC), a machine-learning tool (ViraLM or geNomad), or a homology-based tool (DIAMOND), coupled with Kraken2X for false-positive filtering, maximised recall and F1 score. These performances were maintained despite an increasing rate of induced mutations in Refseq sequences. The homology-based tool, DIAMOND, had the highest CPU and RAM requirements, resulting in the longest runtime for the DVF+DIAMOND combination. In contrast, DVF+DMC was the fastest and least memory-intensive combination, highlighting computational efficiency as an important consideration for clinical and discovery applications.

The application of these combinations on real lung and airborne metagenomic samples have reported a significant additional predicted viral contigs coding for viral proteins. Viral classification and functional compositional analysis indicate that the recovered sequences represent bona fide viral diversity rather than false-positive predictions. They substantially expand the functional characterization of the lung and airway virome beyond existing reference databases. This suggests that the integration of detection strategies, including at least DVF and another tool, may offer a more reliable framework for identifying viral contigs in low-abundance and especially short-read based metagenomic datasets. The consistent presence of DVF in these combinations shows the adaptability of this tool with short viral contigs [18] but with divergent sequences as well.

This combined strategy increased overall sensitivity while providing two solutions to control false positives in viral metagenomic datasets. First, Kraken2X has been reported among the most specific classification tools [23]. It can classify sequences as viral, non-viral, or unassigned. Its large database and strategy of identification based on k-mer homology also allows a larger spectrum of false positive sequences to be identified. Second, CheckV is widely used in virome catalogs for viral sequence classification [13,41]. However, we showed that many viral sequences were passed over CheckV, the majority predicted as undetermined quality. This finding is consistent with previous benchmarking studies [15], which showed that the use of convolutional neural networks based tools improves the performance of viral detection, particularly for CheckV undetermined quality contigs. This is likely because these approaches can capture complex sequence patterns that are difficult to identify using traditional homology-based methods.

Previous prediction tool benchmarking studies have focused on abundant virus ecosystems or *in silico* virus-enriched artificial datasets [31]. However, the performance of these tools on subdominant and divergent viral metagenomes (metaviromes) remains unexplored and is frequently ignored due to small contig sizes. In addition, most benchmark studies evaluate viral detection tools individually to assess their performance [15,31,34]. Nevertheless, the performance of combined tool strategies has rarely been systematically benchmarked [14], although several studies have used unbenchmarked tool combinations to explore and characterize the human gut virome [42–46]. In this study, we showed that viral prediction tools can complement each other by leveraging different algorithms and detection principles.

Our proposed strategy expands the detection capacity of individual viral prediction tools, offering an innovative framework to update the annotation of existing virome catalogs [47]. This multi-algorithmic approach demonstrates robust recovery of short viral contigs (<2000 bp), which are often reported in low-abundance metagenomic dataset.

Combining complementary prediction algorithms can effectively illuminate the "viral dark matter" in undescribed and divergent environmental and host-associated ecosystems [2].

Several limitations should be considered in this study. First, the lack of well-characterized mock communities limited our ability to fully assess the performance of the benchmarked tool combinations. Second, different parameters or analytical approaches may be required for the detection and characterization of DNA and RNA viruses, which warrants further investigation. Notably, several virus prediction tools specifically developed for RNA virus detection were not considered in this study. Third, although short contigs can provide valuable viral information, fragmented sequences may lack complete genomic information and can therefore limit downstream viral characterization. Further studies using well-controlled mock communities and diverse biological datasets are needed to validate and extend these findings.

## Conclusion

This study benchmarks the performance of eleven state-of-the-art bioinformatic tools that can identify viruses, using artificial and real, low-abundant, divergent metagenomic data. Since deep learning tools outperform homolog-based tools, especially on short, low- and undetermined-quality contigs reported by checkV, this strategy provides a promising avenue for viral sequence identification. However, it is limited by a high number of false positives. To address this issue, we performed dual combinations of tools with higher recall capacities and found that DVF+DMC and DVF+ViraLM combinations, coupled with Kraken2X to control false positives, surpassed other single tools and combinations. As previously mentioned, no viral tool is perfect, each having its own strong and weak characteristics [15]. However, we argue that combining tools can increase sensitivity and specificity, especially in divergent sequences with the goal of reducing the viral dark matter in unknown environments.

## List of abbreviations

AI: Artificial intelligence
BLAST: Basic Local Alignment Search Tool
Bp: Base pairs
CNNs: Convolutional neural networks
CT3: Cenote Taker 3
Kb: Kilobase
DMC: DeepMicroClass
DVF: DeepVirFinder
DNA: Deoxyribonucleic acid
FP: False Positive
FN: False Negative
HMM: Hidden Markov Model
LCA: Lowest Common Ancestor
NCBI: National Center for Biotechnology Information
NGS: Next-Generation Sequencing
ORF: Open Reading Frame
PCR: Polymerase Chain Reaction
pVOG: Prokaryotic Virus Orthologous Groups
RefSeq: NCBI Reference Sequence database
RNA: Ribonucleic acid
SARS-CoV-2: Severe Acute Respiratory Syndrome Coronavirus 2
TP: True Positive
TN: True Negative
TNF: Tetranucleotide frequency

## Declarations

### Ethics approval and consent to participate

Not applicable.

### Consent for publication

Not applicable.

### Availability of data and material

All used data were retrieved from the public NCBI reference database and from the published NCBI Bioprojects (PRJNA1110359 and PRJNA854686) previously published. The scripts are available on the GitHub repository: https://github.com/genepii/Virome-Benchmark.

### Authors’ contributions

H.A., and L.J. conceived the study. H.A., S.A., and L.J. drafted the manuscript. S.A. performed bioinformatic analysis. S.A., G.D., M. R., V.N., H.A. and L.J. interpreted results. S.A., G.D., M.R., V.N., L.J and H.A. revised the manuscript. All authors reviewed and approved the final version of the manuscript.

## Acknowledgements

We would like to thank the Hospices Civils de Lyon for using their IT servers at the Hôpital La Croix Rousse and the virome@tlas for insightful discussions.

